# Illuminating the Virosphere’s Dark Matter using Hierarchical Deep Learning

**DOI:** 10.1101/2025.09.22.677955

**Authors:** Chuan Cao, Liang He, Chengping Li, Yuliang Jiang, Chuyue Tang, Chengyue Huang, Yuman Li, Yuan He, Yaosen Min, Haiguang Liu, Tao Qin, Tie-Yan Liu

**Affiliations:** Zhongguancun Academy, China; Zhongguancun Institute of Artificial Intelligence, China; Tsinghua University, China; Peking University, China; Georgia Institute of Technology, United States of America

**Keywords:** Viral dark matter, virus discovery, metagenomics, protein foundation model, open-set recognition, novel viral lineages

## Abstract

Systematic discovery of novel viruses is essential for pandemic preparedness, understanding tumor-associated viruses, developing viral delivery systems, and advancing biomedical applications. Yet, the majority of sequences in metagenomic datasets lack close relatives in existing references, representing a vast viral “dark matter” whose biology and evolution remain largely unknown. The central task is threefold: 1) to determine whether a genome is viral or non-viral, 2) to correctly assign viral genomes to known lineages when possible, and, critically, 3) to recognize when no existing lineage applies and thereby identify candidates for entirely novel viral groups. Existing approaches, which depend on sequence homology or narrow markers, struggle to capture this uncharted viral space. Here we present **DeepVirus**, a hierarchical transformer-based framework that models viral genomes as structured sequences of protein-coding genes. By combining protein-level embeddings from a foundation model with genome-aware representations, DeepVirus not only achieves accurate classification across deep taxonomic hierarchies, but also extends beyond conventional classification to detect and organize candidate novel viral lineages through open-set recognition. Applied to large-scale metagenomic resources, DeepVirus uncovered extensive viral diversity, including previously uncharacterized RNA-dependent RNA polymerases (RdRps), thereby expanding the known evolutionary space of RNA viruses. DeepVirus integrates deep learning with genome-aware open-set discovery to illuminate viral dark matter, providing a foundation for systematic viral taxonomy and advancing exploration of the global virosphere, with broad implications for safeguarding human health.

## 1 Introduction

Viruses are the most abundant biological entities on Earth, shaping ecosystems, driving evolution, and influencing human health [1–6]. They act as vast reservoirs of genetic diversity, regulate microbial communities, and provide indispensable tools for biotechnology and therapy, from viral vectors in gene delivery to oncolytic viruses in cancer treatment. Yet, despite their ubiquity and importance, the majority of viral sequences recovered from metagenomic surveys remain unclassified, constituting a “dark matter” of the virosphere whose roles in biology and evolution are still largely unknown [1, 7–9]. Traditional virus identification strategies, based on sequence homology or narrow functional markers such as RNA-dependent RNA polymerase (RdRp), are inherently limited by reference bias. These methods struggle to detect evolutionarily distant or truly novel lineages, thereby constraining both virus identification and the discovery of entirely new viral categories with coherent functional and evolutionary signatures [7, 8].

Recent advances in AI have begun to address these limitations by introducing alignment-free sequence models and gene-based approaches [10–14]. Alignment-free methods, such as DeepVirFinder [11] and BERTax [12], leverage deep learning to classify viruses directly from nucleotide sequences by identifying informative sequence motifs. Gene-based approaches, exemplified by VirSorter2 [13], rely on curated marker genes or protein profiles to infer taxonomy. While both paradigms have demonstrated strong performance, they remain constrained by database coverage and the scarcity of universal viral markers, particularly for underrepresented groups [7, 15]. Hybrid approaches, such as geNomad [14], attempt to bridge these paradigms, but their effectiveness is still hindered by reference bias and incomplete representation of viral diversity [14, 15]. A promising direction is to harness protein-level information, since protein functional motifs are generally more conserved than primary sequences. Indeed, studies integrating predicted protein structures with sequence information have already revealed previously unrecognized RNA viruses [5, 9]. However, these efforts have largely focused on hallmark genes. Extending such analyses to genome-wide protein repertoires opens the possibility of detecting viruses with low sequence identity but conserved functional signatures [16–19]. Beyond classification, a central challenge remains: discovering viral lineages that fall outside existing taxonomic categories.

To this end, we present **DeepVirus**, a deep learning framework that models viral genomes as structured sequences of protein-coding genes. By integrating protein foundation models with genome-level context, DeepVirus captures functional relationships that go beyond sequence homology and alleviates reliance on narrow marker genes. Critically, DeepVirus incorporates statistical hypothesis testing for open-set recognition [20–22], enabling it not only to classify established lineages but also to detect and organize candidates for novel viral groups. This design provides a scalable, interpretable, and biologically meaningful framework for metagenomic virus exploration. Applied to large-scale metagenomic resources [1–4, 9, 23, 24], DeepVirus illuminates previously hidden branches of the virosphere and creates a systematic resource for viral discovery. Notably, we identify uncharacterized RNA-dependent RNA polymerases (RdRps) that retain hallmark catalytic motifs while displaying lineage-specific divergence, offering preliminary evidence for the existence of novel viral groups and expanding the known evolutionary space of RNA viruses. Future wet-lab validation will be essential to confirm the biological significance of these candidate lineages and to formally integrate them into the broader framework of viral taxonomy.

## 2 Results

### 2.1 DeepVirus is a genome-level framework for illuminating viral dark matter

Accurate virus discovery requires capturing not only the functions of viral proteins but also their organization within a genome. Conventional approaches typically rely on protein homology searches or k-mer statistics, both of which miss novel viruses with limited sequence similarity to known references. To overcome this limitation, we developed DeepVirus, a multi-scale modeling framework that jointly integrates protein-level features with genome-level context.

DeepVirus operates in three stages (Fig. 1). First, each protein sequence within a candidate genome is encoded using a large pre-trained protein foundation model trained with both masked language modeling and pairwise masked language modeling objectives, enabling it to capture local sequence context and residue co-variation. Second, protein embeddings are aggregated in genome order and processed through a self-attention module that models dependencies across the full assembly, generating compact genome-level representations. Finally, these representations feed into a classification head that performs both virus identification (distinguishing viral from non-viral sequences) and hierarchical taxonomy assignment.

**Fig. 1.**
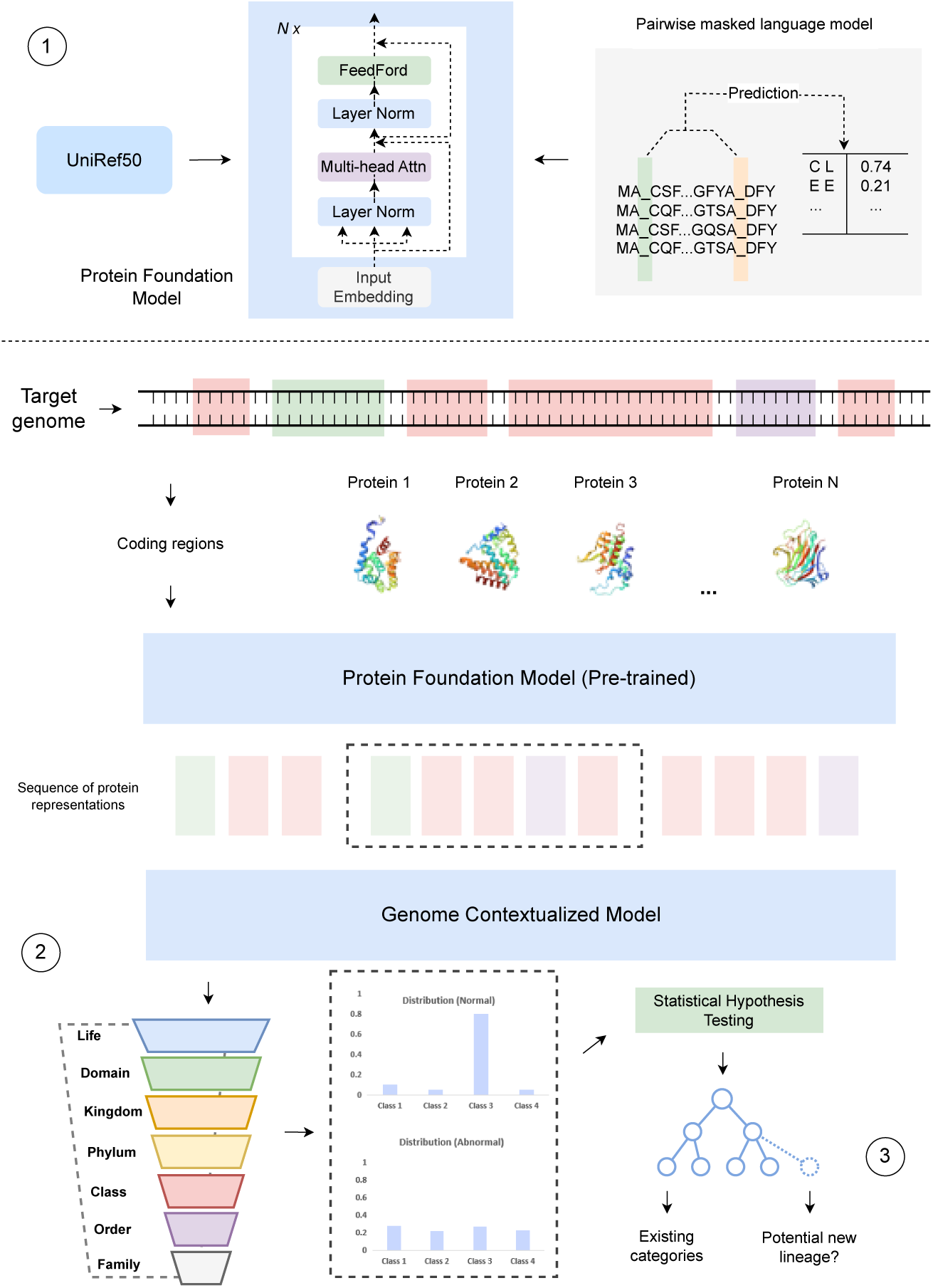
Overview of the DeepVirus framework for viral identification, classification, and novel lineage discovery. DeepVirus integrates protein-level and genome-level information in a three-stage process. (1) Protein encoding: Viral contigs are parsed into coding regions, and each protein is embedded using a pre-trained protein foundation model, capturing both local features and genome-wide organization. (2) Hierarchical classification: The ordered protein representations are processed by a transformer encoder with a hierarchical classification head, enabling accurate predictions across taxonomic ranks while enforcing consistency between levels (e.g., order, family). (3) Novel lineage discovery: Sequences that cannot be confidently assigned to known categories are identified through open-set recognition and clustered into coherent groups, supporting the systematic discovery of previously unrecognized viral lineages.

For the foundation model pretraining, we adopted a pairwise masked language modeling objective [25, 26], which explicitly encourages the model to learn functional interactions beyond conventional masked language modeling. To train the classification module, we curated genome assemblies from NCBI RefSeq [27], splitting them chronologically to prevent data leakage and reserving later-released genomes as the held-out test set. Proteins were either taken directly from RefSeq annotations or predicted de novo with Prodigal [28], ensuring robust evaluation across both curated and real-world data.

To identify novel viral lineages, we estimated the distribution of model logits for each taxonomic category and applied Gaussian hypothesis testing. Genomes confidently assigned to an existing distribution were classified accordingly, while those rejected across all categories were flagged as candidates for new viral groups.

This design allows DeepVirus to leverage high-quality protein embeddings while integrating genome-wide organizational signals. By learning jointly from these two levels, DeepVirus provides a powerful framework for discovering and classifying viruses, including those highly divergent from existing references. In doing so, it lays the foundation for systematically charting the vast reservoir of viral dark matter in ecosystems ranging from ocean microbiomes to human-associated viromes.

### 2.2 DeepVirus accurately identifies and classifies viruses

We first evaluated DeepVirus on RefSeq genomes released after December 31, 2022, yielding 28,702 assemblies (14,820 viral and 13,882 non-viral) comprising 100 million proteins. This chronological split ensured strict separation between training and test data. For comparison, we benchmarked DeepVirus against MMseqs2 (homology-based), BERTax (DNA-level transformer), DeepVirFinder (k-mer–based deep learning), and geNomad (hybrid alignment-free and gene-based). Predictions from baseline tools were aggregated to the genome level via majority vote.

In the binary virus identification task (Fig. 2), DeepVirus achieved near-perfect performance, with accuracy of 0.999 and MCC of 0.997, substantially surpassing all baselines. These results highlight the importance of combining protein-level functional signals with genome-level organization, enabling sensitive and specific detection of divergent viruses that evade traditional homology or k-mer–based heuristics. From a biological standpoint, this capability ensures that novel viruses can be confidently distinguished from host or microbial contamination, reducing false positives that often confound virome studies.

**Fig. 2.**
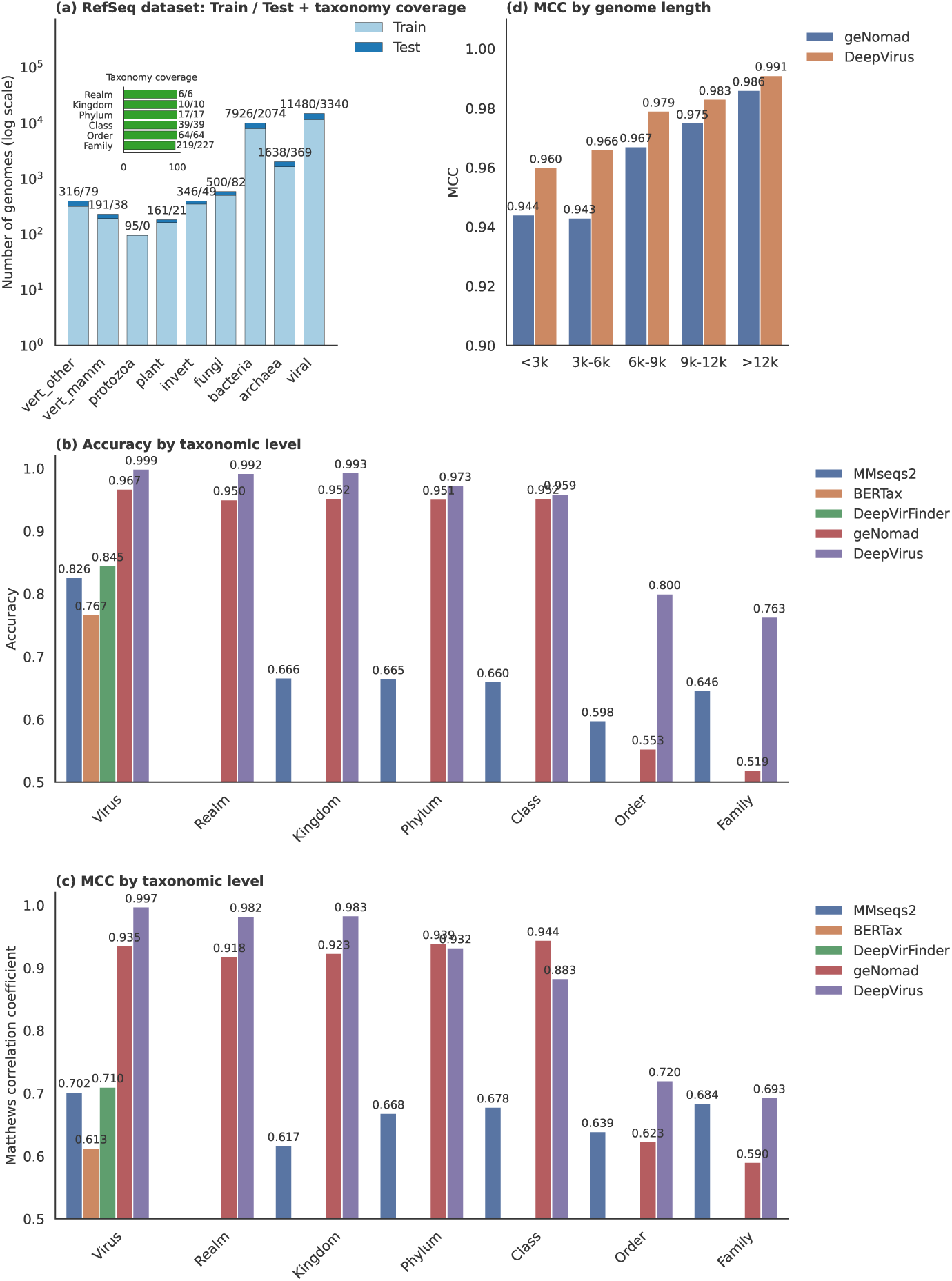
Dataset composition and classification performance of DeepVirus. (a) Distribution of training (light) and test (dark) genomes across major host categories. Inner vertical bars indicate the taxonomic breadth of the dataset, illustrating coverage across hierarchical levels. (b) Comparative classification accuracy of DeepVirus and baseline methods across taxonomic ranks, showing consistent improvements. (c) Comparative Matthews correlation coefficient (MCC) of DeepVirus versus baselines, further highlighting robust gains across ranks. (d) Head-to-head evaluation of DeepVirus and geNomad on the geNomad dataset, stratified by genome size, demonstrating superior performance of DeepVirus across diverse genome lengths.

When restricted to viral genomes, DeepVirus also outperformed baselines across six hierarchical taxonomic levels. At higher ranks (realm, kingdom, phylum), DeepVirus achieved accuracies above 0.97 and MCCs above 0.93. Its advantage was even more pronounced at finer ranks such as order and family, where accurate classification is critical for linking new viruses to host range, replication strategy, and ecological function. Thus, DeepVirus not only identifies viruses but also provides biologically meaningful annotations that accelerate downstream hypothesis generation.

We further benchmarked DeepVirus against geNomad on its evaluation datasets stratified by genome length (Fig. 2). DeepVirus consistently out-performed geNomad across all bins, with particularly strong gains for short genomes (*<*3 kb), where MCC increased from 0.944 to 0.960. This robustness is especially relevant for metagenomic studies, where viral contigs are often fragmented or incomplete. DeepVirus’s ability to integrate sparse protein evidence into coherent genome-level predictions greatly improves recovery of viral diversity from environmental data.

### 2.3 DeepVirus can discover new virus lineages beyond known classification

To test DeepVirus’s capacity for detecting novel lineages, we applied statistical hypothesis testing to phylum-level distributions of RefSeq genomes. Each newly predicted viral genome was compared against all known phyla via Z-tests under a normality assumption. Genomes classified as viral but rejected across all phyla were flagged as candidates for novel groups (see Methods).

We further conducted leave-one-phylum-out experiments to rigorously assess generalization. For each phylum with more than 100 genomes, the model was trained without that phylum and then applied to its withheld genomes. A second round of discovery was performed by assigning new labels to rejected genomes, retraining the model, and re-evaluating remaining data, simulating the iterative process of uncovering an unseen lineage. Results are summarized in Fig. 3.

**Fig. 3.**
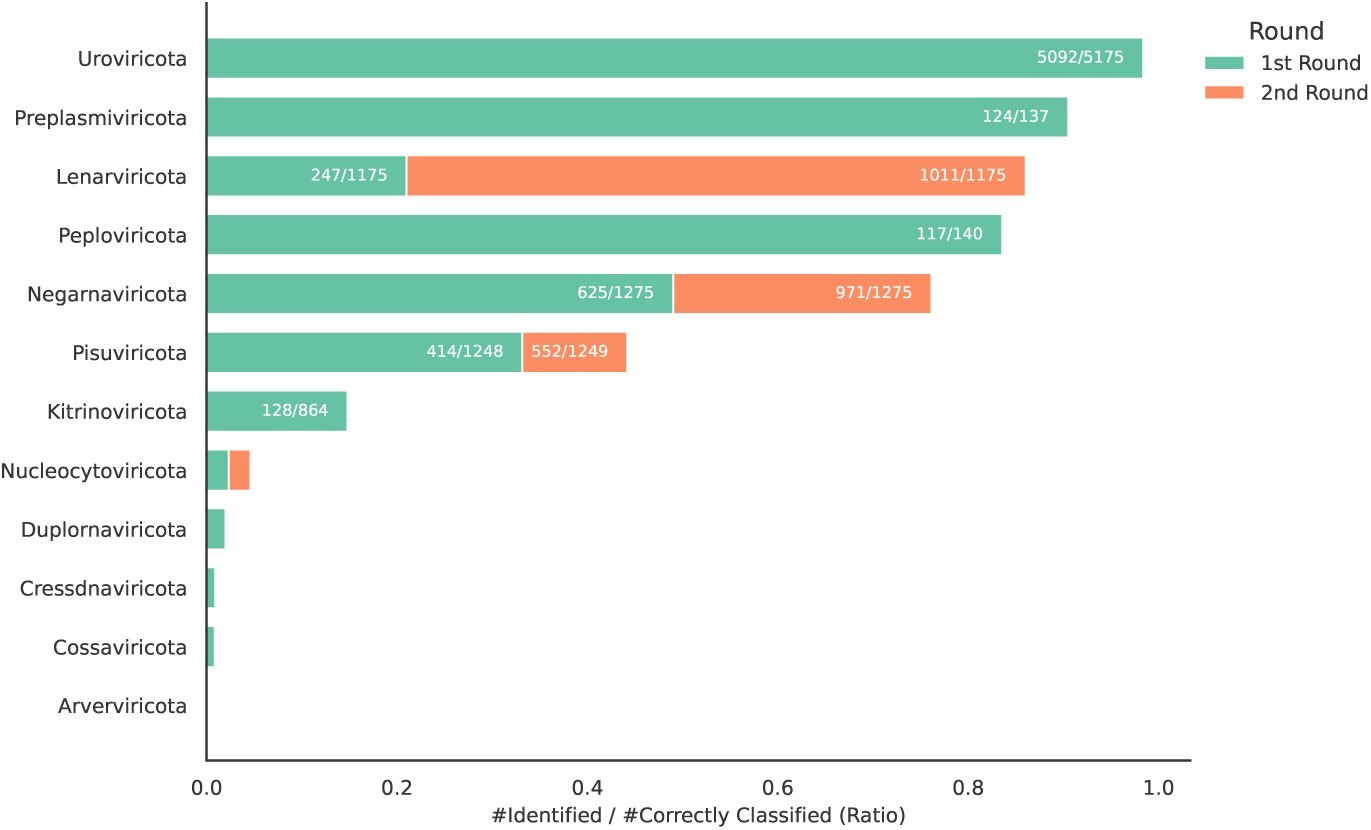
Leave-one-phylum-out evaluation of DeepVirus for re-discovery of viral lineages. Shown is the fraction of genomes correctly identified by DeepVirus when entire viral phylum genomes are withheld during training. Bars represent identification rates from the first and, where applicable, second rounds of re-training and classification, demonstrating the model’s ability to detect novel phyla with high specificity. Notably, approximately half of the withheld phyla can be effectively re-discovered with high recall, underscoring DeepVirus’s capacity to recover broad evolutionary groups absent from training data.

DeepVirus demonstrated strong capacity to rediscover known phyla. In the first round, *Preplasmiviricota, Peploviricota*, and *Uroviricota* were almost completely recovered, while *Lenarviricota, Negarnaviricota*, and *Pisuviricota* showed moderate recovery that improved substantially after retraining. Together, these results indicate that DeepVirus can effectively reconstruct roughly half of the tested phyla. For phyla with fewer samples or closer similarity to others, a smaller fraction of genomes was flagged as distinct, reflecting the conservative but reliable nature of the statistical framework.

### 2.4 DeepVirus reveals the viral dark matter in massive real-world metagenomic data

Given its strong performance on curated data, we next applied DeepVirus to massive metagenomic datasets. Viruses dominate marine ecosystems and contribute substantially to microbial community dynamics and global biogeochemical cycles [29]. Giant viruses of the phylum *Nucleocytoviricota*, in particular, are abundant and diverse, yet remain incompletely characterized [30]. Despite advances, scalable genome-level virus identification frameworks for metagenomic data remain scarce.

We first analyzed a large ocean metatranscriptome dataset [31]. After filtering assemblies with fewer than four proteins or proteins shorter than 20 amino acids, the dataset comprised 1,943,085 genomes encoding 10 million proteins. DeepVirus identified 14.2% of genomes as high-confidence viral (*p* ≥ 0.99), 92% of which overlapped with annotations from the original study. Remarkably, nearly 12% of these high-confidence calls represented potentially novel lineages absent from known phyla (Fig. 4).

**Fig. 4.**
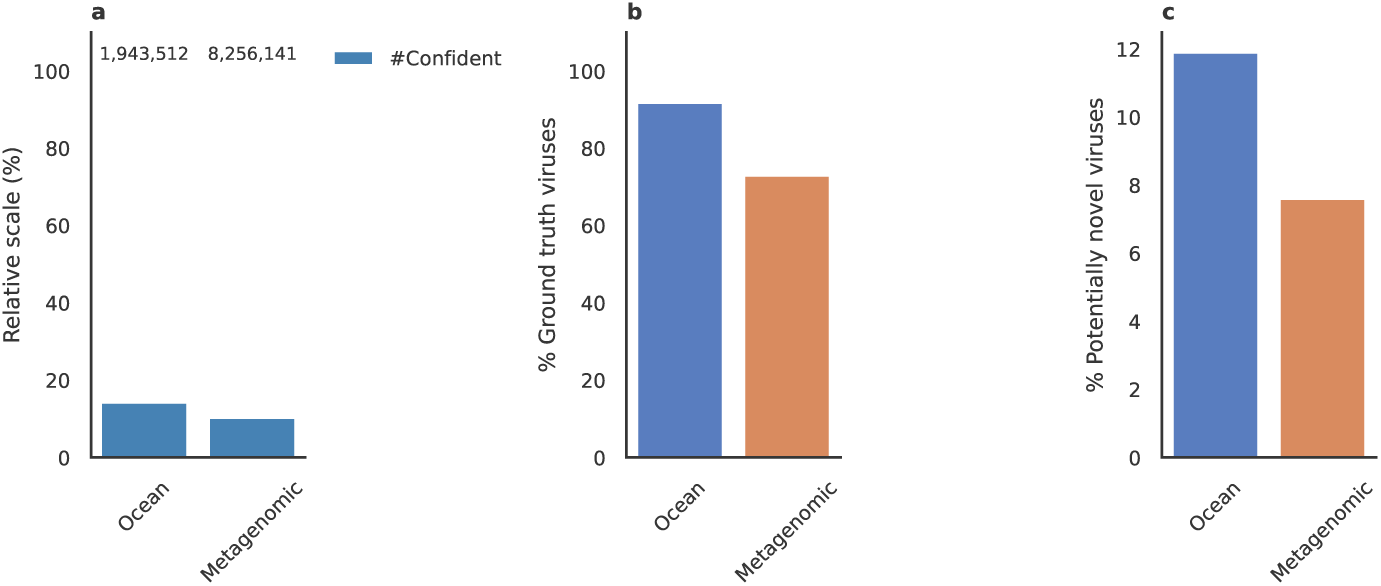
Viral discovery in ocean and metagenomic datasets using DeepVirus. a, Proportion of high-confidence predictions (*p* ≥ 0.99) among all genomes. b, Evaluation of prediction quality, showing the percentage of ground-truth viruses recovered (% of gt) of the tested genomes. c, The proportion of potentially novel viruses identified (% potential novel). DeepVirus demonstrates robust detection across diverse datasets, capturing both known and previously uncharacterized viral sequences.

We then applied DeepVirus to an even larger metagenomic dataset comprising 8,256,141 assemblies and more than 900 million proteins [9]. DeepVirus classified 10.2% of genomes as high-confidence viral, with 72.9% overlapping previous annotations. Notably, 7.6% of predictions represented candidate novel lineages, underscoring the ability of DeepVirus to reveal hidden viral diversity at scale.

### 2.5 Discovery and validation of novel RdRps from DeepVirus-identified viruses

RNA-dependent RNA polymerase (RdRp) is one of the most conserved proteins across RNA viruses, universally required for replication and transcription [32]. Its evolutionary conservation and hallmark sequence motifs make it a cornerstone for RNA virus discovery and classification [33].

To evaluate whether DeepVirus uncovers genuinely novel RNA viruses, we focused on candidate genomes meeting two stringent criteria: absence of similarity to cellular proteins and presence of conserved RdRp motifs. Most identified RdRps retained the hallmark catalytic motif C (GDD) [34], supporting their assignment as viral polymerases (Fig. 5a). Phylogenetic analysis revealed that many clustered into clades distinct from known families, highlighting previously unrecognized lineages and expanding the evolutionary breadth of RNA viruses (Fig. 5b).

**Fig. 5.**
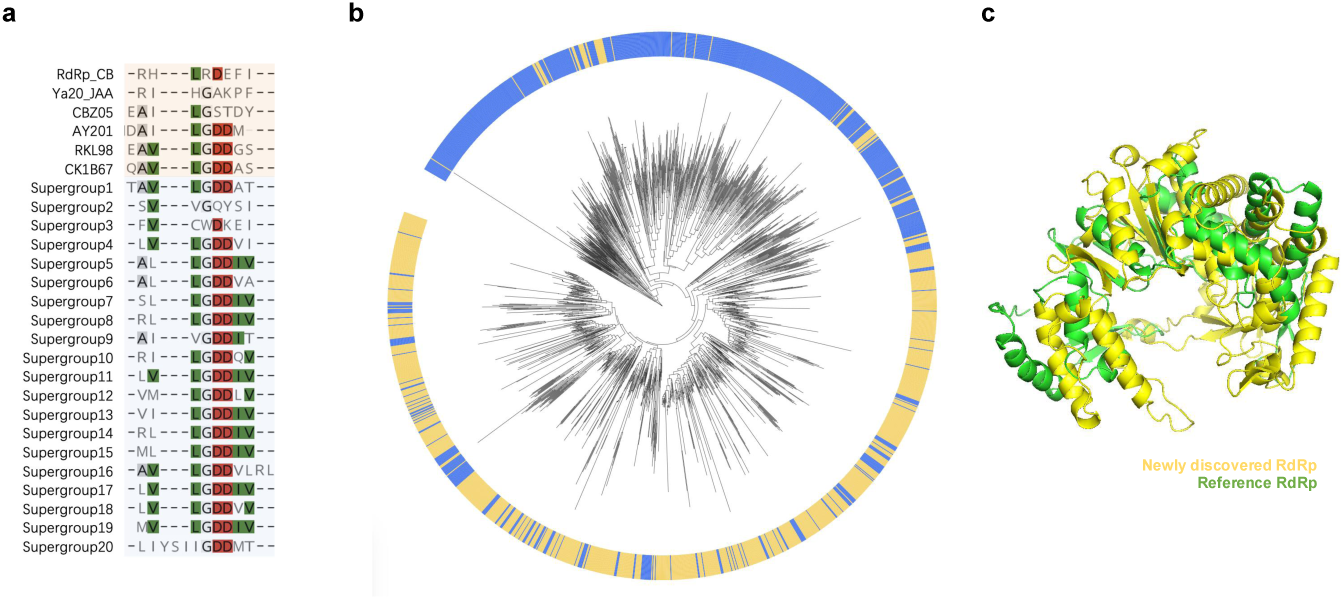
Discovery and validation of novel RdRps identified by DeepVirus. a, Multiple sequence alignment of representative candidate and reference RdRps highlighting conservation of the catalytic motif C (GDD). b, Maximum-likelihood phylogeny of RdRps constructed with IQ-TREE3, showing newly discovered RdRps forming distinct clades (yellow) separate from reference lineages (blue). c, Predicted structure of a representative candidate RdRp (yellow) superimposed on a reference viral RdRp (green), both adopting the canonical right-hand fold; the alignment yielded an overall RMSD of 8.4 Å.

Structural modeling further reinforced these findings. Newly identified RdRps consistently adopted the canonical “right-hand” architecture of palm, fingers, and thumb domains, with strong conservation of the catalytic palm subdomain harboring the GDD motif. At the same time, substantial divergence was observed in peripheral regions, particularly in the fingers and thumb domains, reducing overall similarity to known RdRps. This combination of conserved catalytic cores with lineage-specific peripheral variation suggests the presence of evolutionarily novel polymerases (Fig. 5c).

Together, motif alignments, phylogenetic relationships, and structural comparisons provide convergent evidence that DeepVirus uncovers previously uncharacterized RdRps, supporting the discovery of new RNA viral lineages and extending the known evolutionary landscape of RNA viruses.

## 3 Discussion

Our findings demonstrate that hierarchical modeling of viral genomes not only achieves accurate classification across diverse taxonomic levels but also establishes a scalable framework for the systematic discovery of previously unrecognized viral lineages. By moving beyond conventional homology- and marker-dependent strategies, this approach substantially broadens the reach of computational methods, enabling them to capture both the breadth and depth of viral diversity. In doing so, it lays a foundation for more comprehensive and systematic mapping of the global virosphere.

Applying this framework to large-scale metagenomic resources, we uncovered vast regions of viral “dark matter” that have remained inaccessible to traditional methods. These results highlight the power of combining advanced deep learning with rapidly expanding genomic datasets to accelerate virus discovery at an unprecedented scale. The catalog of candidate viruses generated here provides a valuable community resource, and we invite collaborative efforts to investigate the ecological, evolutionary, and biomedical implications of this newly revealed diversity.

While our computational predictions are supported by extensive in-silico analyses, the functional and biological relevance of these putative viral groups remains to be confirmed. Experimental validation, through viral isolation, host identification, and structural and functional characterization, will be critical to place these candidate lineages within a broader biological and taxonomic context. Looking ahead, the integration of experimental virology with computational discovery pipelines will be essential to fully realize the potential of this approach, deepening our understanding of viral evolution and advancing applications in health, ecology, and biotechnology.

## 4 Methods

Our proposed DeepVirus is a genome modeling framework for virus identification and taxonomic classification. It has two components: (i) hierarchical modeling of genome sequences for multi-level classification, and (ii) discovery of novel virus lineages through open-set recognition built on the trained genome classifier. The core idea of the genome modeling module is to represent each genome through its encoded proteins, based on the biological observation that microbial genomes are highly compact in coding regions, where functional interactions among DNA elements are concentrated. To achieve this, DeepVirus employs a pretrained protein language model to generate protein representations, which are then processed by a Transformer trained on hierarchical genome data. For open-set recognition, statistical hypothesis testing is applied to the classification outputs. If a genome is confidently identified as viral but rejected by all known categories, this serves as evidence that it may belong to a previously uncharacterized viral lineage.

### 4.1 Protein Encoder Pretraining

We pre-trained a protein language model to obtain sequence embeddings that capture structural and functional dependencies among residues. The model follows the masked language modeling (MLM) paradigm [35, 36], where a subset of residues (typically 15%) is masked and the encoder is trained to recover them from surrounding context by minimizing the negative log-likelihood. Unlike natural language, however, protein sequences encode three-dimensional structures and functions through inter-residue dependencies, which are not well captured by token-wise predictions alone. To address this, we adopted the model from the recent work which extends the standard MLM objective with a pairwise masked language modeling (PMLM) task [25, 26], in which the model jointly predicts pairs of masked residues. This formulation accounts for co-evolutionary signals by recognizing that the joint probability of two residues cannot be factorized into independent predictions. The final training loss is simply the sum of the standard MLM loss and the PMLM loss, allowing the encoder to learn both local sequence context and global co-variation patterns. The encoder is implemented as a multi-layer Transformer with standard self-attention and feed-forward blocks [37] For the input protein sequence *s* = [*s*_*cls*_, *s*_1_, *s*_2_, …, *s*_*k*_, *s*_*eos*_], the one-hot embedding and positional embedding are applied:

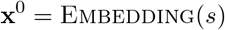

following by the applied Transformer encoder layers as (for *l* ∈ [1‥*L*]):

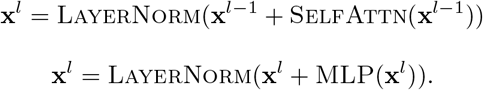

Pretraining was conducted on the UR50 dataset (about 30 million protein sequences) using the BERT masking strategy [35] (80% replaced by a mask token, 10% by a random residue, 10% unchanged) for the two prediction heads for residue-wise and pairwise tasks. The resulting pretrained encoder yields rich contextual embeddings for proteins, capturing both local residue features and higher-order co-variation signals. These embeddings serve as the foundation for downstream genome modeling, where proteins are treated as compositional units analogous to words in a document, and their arrangement is exploited to infer genome-level properties.

### 4.2 Genome Multi-level Classification

For the input sequence of protein sequence representation 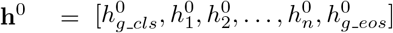, where 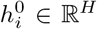 is the vectorized representation for the *i*-th protein sequence (i.e., 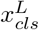, the output hidden vector for the CLS token of the protein) and *i* ∈ [1‥*n*], 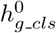 and 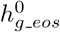 are manually added vec-tors as the CLS token and the EOS token for the whole genome representation. For the input embeddings, we apply iteratively *K* Transformer encoder layers to the initial or intermediate representations:

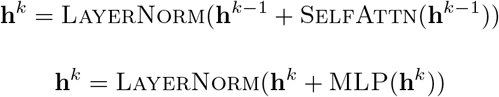

Viral taxonomy is inherently hierarchical, spanning multiple ranks such as family, genus, and species. To reflect this structure, we design a hierarchical classification head that jointly models global dependencies across the taxonomy and local specialization at each level. The architecture follows the principle of hierarchical multi-class classification [38], where predictions are refined through a recurrent flow of information.

Let *h* ∈ ℝ^*H*^ denote the pooled genome representation (genome CLS token) from the genome model output, i.e., 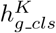. After dropout and a non-linear transformation, we initialize the global flow with *h*:

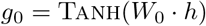

At each taxonomy level *m* in the recurrent hierarchy, the state is updated by concatenating the previous hidden state *g*_*m−*1_ with the original feature representation *h*:

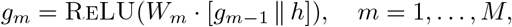

where *m* indexes the hierarchy levels and *M* is the total number of levels. In parallel, each hidden state *g*_*m*_ is transformed into a prediction through a two-layer feed-forward network, followed by a softmax function to produce the probability distribution over the categories:

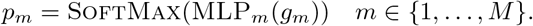

### 4.3 Loss Function

We use the standard cross-entropy loss ℒ_classifier_ for classification accuracy at each taxonomy level. In addition, viral taxonomy follows a tree-structured hierarchy, where predictions at finer levels (e.g., genus or species) must remain consistent with broader assignments (e.g., family or order). To enforce this property, we incorporate a hierarchical violation loss inspired by [38]. Without such a constraint, the model could assign a high probability to a parent class (e.g., a realm) while simultaneously predicting a child class (e.g., a kingdom) that does not belong to it. Formally, a violation occurs when the predicted probability of a parent node *p*_parent_ is smaller than the sum of the predicted scores of its child nodes *p*_child_. The hierarchical violation loss is defined as:

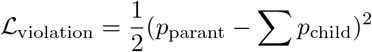

The final training objective combines the two terms:

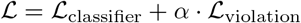

where *α* ≥ 0 controls the strength of the hierarchical consistency regularization. Because the magnitude of ℒ_violation_ is relatively small, we set *α* = 100 to scale its contribution to a level comparable with the classification loss.

Viral taxonomy classification is highly imbalanced, with some lineages heavily represented in reference databases and others represented by only a handful of genomes. Without addressing this imbalance, models risk being biased toward majority classes while neglecting rare but biologically meaningful taxa. In the RefSeq database, for example, our training set of 11,480 viral genomes includes 4,480 genomes from kingdom Heunggongvirae, but only 6 from kingdom Helvetiavirae. At finer levels such as order and family, many classes contain only one or two training samples, in stark contrast to other classes that include hundreds. To mitigate this imbalance, we re-weight the classification loss based on the training sample distribution. Intuitively, mis-classifications of minority classes are penalized more heavily than those of majority classes, encouraging the model to learn stronger representations for rare taxa. Formally, let *N*_*c*_, *c* ∈ *C* denote the number of training samples for class *c*. The weight for class *c* is defined as:

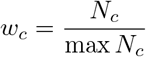

In the classification loss, a single sample’s loss is scaled by the weight of its label class. In practice, models trained with loss re-weighting show consistent improvements, particularly at lower taxonomy ranks, across multiple model architectures. Importantly, this adjustment improves the model’s ability to capture rare taxa, which are often of greatest interest for detecting novel or emerging viral lineages.

### 4.4 Training

We trained our genome-level virus classification model using a distributed setup with 8 GPUs via PyTorch. The genome encoder follows a BERT-style architecture with a hidden size of 1,280 and 8 Transformer layers. Training was performed for 300 epochs with a batch size of 64. The optimizer was Adam with a learning rate of 1 × 10^*−*5^, *β*_1_ = 0.9, *β*_2_ = 0.99 and no weight decay. To stabilize training, gradient clipping with a maximum norm of 1.0 was applied. The taxonomy hierarchy spans 7 levels. Because the distribution of viral taxa is highly imbalanced, we applied classification loss re-weighting to upweight minority classes. Additionally, the hierarchical consistency loss was used to enforce taxonomic validity across levels. For efficiency, genome lengths were divided into 100 bins and sampled in a partially randomized manner at each epoch. Models were trained with both taxonomy-aware loss reweighting and hierarchical consistency regularization enabled.

### 4.5 Datasets

The training dataset consisted of viral genomes from RefSeq, along with non-viral genomes (archaea, bacteria, fungi, protozoa, vertebrate mammalian, vertebrate other, invertebrate, and plant) included as negative samples to improve discrimination. For model training and evaluation, we curated viral genome sequences from the RefSeq database. To prevent information leakage, the dataset was split by release date: all genomes released on or before 2022-12-31 were used for training and validation, while genomes released afterward were held out as the test set. Within the training set, sequences were further split into 80% for training and 20% for validation. Protein-coding regions within each genome were annotated by the dataset itself. These protein sequences were encoded using the pairwise masked language model described above, generating compact embeddings that capture residue-level co-variation and structural information. These embeddings were then used as input to the genome-level classification model. To assess real-world generalization, we collected independent viral genome sequences from marine environments (Ocean data) and other publicly available whole-genome datasets. Since protein annotations were not always available for these genomes, we predicted coding sequences using Prodigal, and these predicted proteins were subsequently encoded with the same model before classification. The RefSeq dataset contains 11,480 viral genomes in the training set and 3,340 viral genomes in the test set, covering 7 taxonomic levels from realm to family. On average, each genome contains about 45 protein sequences, though this varies by virus family. The number of classes at each taxonomic rank is highly imbalanced: for example, the kingdom level includes 10 major classes, while lower levels such as family contains more two hundred classes, many with only a few genomes. This highlights the importance of hierarchical classification and loss re-weighting to handle rare taxa effectively.

### 4.6 Baselines and Evaluation Protocol

To evaluate the performance of our genome-level classification framework, we conducted systematic experiments on viral genomes held out from training. Metrics were computed at multiple taxonomic ranks, including realm, kingdom, phylum, class, order, family, and genus, to assess both coarse- and fine-grained classification. We report accuracy and MCC score at each level. For hierarchical evaluation, predictions were constrained to satisfy taxonomic consistency, and misclassifications that violate the hierarchy were separately tracked to assess the impact of hierarchical consistency loss. We compared our method against four state-of-the-art baselines for viral genome classification and detection:

- MMseqs2 - A sequence similarity-based method using sensitive alignment for taxonomic assignment. We used default parameters recommended for viral genome classification, and predictions were aggregated at each taxonomy level according to the best-hit criteria.
- BERTax - A protein language model-based virus classifier that encodes genome-derived protein sequences using a pretrained Transformer model. We followed the authors’ recommended training and evaluation protocol, ensuring fair comparison on the same test set.
- DeepVirFinder - A deep learning-based viral contig classifier that predicts viral sequences from metagenomic fragments. For genome-level evaluation, individual contigs were aggregated, and genome-level labels were assigned based on the majority vote of contig predictions.
- geNomad - A hybrid tool for virus discovery and taxonomic annotation. Predictions were collected using default model parameters, and taxonomy labels were mapped to the corresponding hierarchical levels to allow direct comparison.

All methods were evaluated on the same held-out RefSeq viral and non-viral genome dataset. For methods that do not explicitly support multi-level taxonomy prediction, outputs were mapped to the corresponding ranks using NCBI taxonomy annotations.

### 4.7 New Virus Lineage Discovery

To identify potential novel viral lineages, we applied a statistical hypothesis testing procedure for open-set recognition to the predicted probabilities from our genome classification model. This approach evaluates whether the predicted probability for a genome belonging to any known class is statistically consistent with the distribution of probabilities observed for that class in the training set. Specifically, for each taxonomic rank (e.g., phylum), we first transformed the predicted probabilities *p* using the logit function:

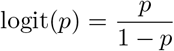

to stabilize variance and avoid boundary effects. For each class, the mean and standard deviation of logit-transformed probabilities were estimated from genomes confidently assigned to that class (probability ≥ 0.99). For a query genome, the null hypothesis assumes that its predicted probability for a given class comes from the corresponding class distribution. A one-sided normal test was performed for each class by computing the z-score of the genome’s logit probability relative to the class mean and standard deviation:

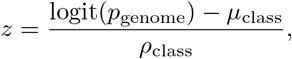

where *µ*_class_ and *ρ*_class_ are estimated from the logits of the training data for each category. If the z-score falls within the acceptance region (p-value *>* 0.05), the genome is considered statistically consistent with the class. If none of the classes at a given taxonomic rank pass the test, the genome is labeled as a rejected candidate, indicating it may belong to a novel lineage. This procedure was applied across all relevant taxonomic ranks, and genomes flagged as rejected were exported for further analysis. By incorporating statistical hypothesis testing on top of the classification probabilities, the model can systematically identify genomes that deviate significantly from known classes, supporting open-set recognition of previously unobserved viral lineages. To further enhance discovery, we introduced a second round: genomes identified as potential new lineages in the first round were assigned an additional category label and incorporated into the training set. The model was then retrained and reapplied to the remaining data, enabling iterative expansion of recognized viral groups.

### 4.8 Viral protein prediction and RdRp identification

Viral genomes identified as putative new lineages by DeepVirus were based on open reading frame (ORF) prediction using Prodigal v2.6.3 [28]. Predicted protein sequences were functionally annotated with eggNOG-mapper v2.1.13 [39], facilitating the detection of proteins associated with RNA-dependent RNA polymerase (RdRp) activity. Candidate RdRp sequences were extracted based on annotation results and cross-referenced with curated datasets from RdRp-Scan [40] and relevant literature to establish a comprehensive reference set.

Candidate and reference RdRp sequences were aligned using MAFFT v7.525 [41]. Conserved catalytic motifs, including motif A (DxD), motif B (SGxxxT), and motif C (GDD) [34], were examined to confirm the presence and integrity of active sites. Phylogenetic reconstruction was performed with IQ-TREE v3.0.1 [42] using 1,000 ultrafast bootstrap replicates, enabling evolutionary placement of candidates and the identification of potential novel clades.

Representative RdRp candidates from putative novel clades were structurally modeled using AlphaFold3 [43]. Predicted structures were examined for the canonical “right-hand” polymerase fold, consisting of palm, fingers, and thumb domains, with emphasis on the spatial organization of the catalytic GDD motif within the palm subdomain. Structural comparison and visualization were carried out in PyMOL v2.6 [44], allowing identification of conserved catalytic features as well as lineage-specific structural deviations relative to reference RdRps.

